# Brain-wide cell-type-specific noradrenergic modulation of the transcriptome

**DOI:** 10.1101/2025.05.09.653136

**Authors:** Cody Slater, Timothy Lantin, Steven M. Wellman, Litian Jia, Gabriel Gonzalez, José L. McFaline-Figueroa, Qi Wang

## Abstract

Neuromodulatory systems such as the locus coeruleus–norepinephrine (LC-NE) system exert a widespread influence on brain function, yet the transcriptional consequences of such neuromodulatory perturbations remain largely unknown across the many unique cell types in the brain. In this study, we establish a generalizable framework to map brain-wide, cell-type-specific gene expression changes in mice following *in vivo* chemogenetic activation or inhibition of LC neurons. Single-nucleus RNA sequencing revealed that LC perturbation induces widespread but highly cell type- and region-specific transcriptional program changes, shaped by the distribution of adrenergic receptor subtypes. These findings support a model in which a shared global signal of neuromodulatory tone can produce discrete, context-dependent cellular outcomes through distinct molecular gating mechanisms of cell-type-specific adrenergic receptor subtype combinations. By establishing gene expression as a quantifiable metric of neuromodulatory control, this study lays the foundation for transcriptionally informed interventions capable of modulating brain functions with cellular precision.

## Introduction

Neuromodulators play a critical role in mediating the flexibility of neural information processing underlying various perceptual and cognitive functions [1, 2, 3, 4, 5, 6, 7, 8, 9, 10, 11]. Numerous molecules, including biogenic amines, neuropeptides, cytokines, and gases, act in concert to dynamically modify circuit output [3, 12, 8, 13, 14, 15]. The result is a functional reconfiguration of otherwise hard-wired networks through the alteration of properties within individual neurons, on synapses, and across the local microenvironment [3, 16, 17, 18, 19, 20, 21]. Most neuromodulatory systems use a volumetric mode of signaling instead of direct synaptic transmission, facilitating simultaneous influence over a broad population of cells [8, 18, 9, 22, 23]. This mechanism affects control over global patterns of brain activity and plays a critical role in switching between distinct brain states (e.g., sleep vs. arousal) and the transition between them [5, 24, 25, 26, 27]. A comprehensive understanding of the role neuromodulation plays in healthy brain function is far from complete and is impeded by the complexity of how these molecules shape neural activity.

In mammals, the ascending arousal system comprises the most well-characterized set of neuromodulators and includes the noradrenergic, serotonergic, dopaminergic, and cholinergic systems [28, 8]. The noradrenergic system is of particular interest due to its largely singular source, from one of the smallest but most extensively projecting nuclei in the brain. The primary source of norepinephrine (NE) is the locus coeruleus (LC), which is a small, bilateral pontine nucleus. Its axons project broadly throughout the forebrain, midbrain, and hindbrain, exerting widespread influence over arousal, learning, and sensory processing. [4, 28, 16, 29, 30, 11]. Individual LC-NE neurons receive and integrate information from many brain regions and, likewise, broadcast widely across mostly non-specific projections [16]. As such, each LC-NE neuron integrates many inputs into a signal that is distributed widely throughout the brain. This input-output relationship is largely homogeneous, though there is evidence of some additional sub-circuit specificity, such as medulla-projecting LC-NE neurons receiving disproportionately less input from the central amygdala [16]. Others have described a limited modularity of the LC, including that individual LC neurons collateralize to innervate functionally related circuits such as somatosensory thalamus and somatosensory cortex [31].

The extent to which LC-NE sub-circuits can facilitate heterogeneous output remains insufficient to explain the full scope of context-dependent NE modulation. This is due, in part, to the diverse array of cell types with unique molecular and functional properties [32]. The noradrenergic system exerts its influence through a differentially expressed array of competing receptor subtypes that alter the intrinsic or extrinsic properties of a defined neural circuit in a region-specific manner [18]. These NE-mediated changes can be rapid and reversible or long-lasting and persistent, depending on the context and duration of signaling. In this setting, cell-type-specific adrenergic receptor (AR) subtypes play an important role in receiving and decoding global LC-NE signaling in a region- and circuit-specific manner. ARs are G protein-coupled receptors (GPCRs) divided into two main classes: alpha-adrenergic receptors (*α* ARs) and beta-adrenergic receptors (*β* ARs). Each class is further divided into subtypes, with *α* ARs including *α*_1*a*_, *α*_1*b*_, *α*_1*d*_, *α*_2*a*_, *α*_2*b*_, and *α*_2*c*_, and *β* ARs including *β*_1_, *β*_2_, and *β*_3_ [33, 34, 35]. *α*_1_ ARs activate the *P LC*-*IP*_3_-*DAG* signaling pathway through a *G*_*q/*11_ protein subunit, leading to increased intracellular calcium levels, activation of protein kinase C (PKC), and regulation of calcium-sensitive processes [36, 37]. *α*_2_ ARs inhibit adenylate cyclase through a *G*_*i*_ subunit, decreasing cyclic adenosine monophosphate (cAMP) levels and protein kinase A (PKA), as well as activating *K*^+^ and inhibiting L- and N-type *Ca*^2^+ channels through the *βγ* and *G*_*O*_ subunits, respectively, of the *G*_*i*_ GPCR [38, 37]. In direct contrast, all *β* ARs act through a *G*_*s*_ GPCR subunit to activate adenylate cyclase, increase cAMP levels and promote neurotransmitter release [39, 40, 37]. The relative affinities of each subtype for NE is estimated to be *α*_2_ > *α*_1_ » *β*_1_ > *β*_2_ > *β*_3_, with *α*_1_ ARs showing the strongest localization to specifically post-synaptic neurons [41, 42, 43].

The recent completion of several brain-wide transcriptomic atlases, such as the Allen Brain Cell (ABC) atlas [44], provides an unprecedented view of the molecular identity of individual cell types across functional subdivisions of the mouse brain [45, 46, 47, 44]. The result is a high-resolution description of each transcriptionally distinct cell type, including cell-type-specific expression of certain receptors, such as ARs. These molecular atlases offer a new opportunity to observe a global view of neuromodulation. Volumetric release of NE exerts direct input on cells that express ARs within local circuits. The result of that input is determined by the functional properties of the receiving cell type and communication within the local network. There is now a substantial need to connect these descriptive atlases with methods to measure the impact on each cell type in a global manner. Recent advances in single-cell and single-nucleus RNA sequencing (snRNA-seq) and *in vivo* chemogenetic manipulation provide a powerful platform for mapping the transcriptional consequences of neuromodulatory signaling across the whole brain in a single experiment.

To obtain a systematic understanding of how discrete modes of noradrenergic activation or inactivation affects the brain wide molecular states across diverse cell populations, we have developed a generalizable transcriptomic framework for profiling cell type- and region-specific gene expression changes following defined neuromodulatory perturbations. As a proof-of-principle, the murine LC-NE system was activated or inactivated *in vivo* using DREADDs (Designer Receptors Exclusively Activated by Designer Drugs), and neural tissue sampled from across the brain was profiled using snRNA seq. Our data offer molecular insights into how the LC exerts its broad physiological effect and demonstrates the utility of using transcriptomic readouts in intact, perturbed neural circuits to reveal the global influence of neurochemicals in the brain. By establishing gene expression as a measurable and controllable substrate for neural manipulation, this work lays a foundation for neuromodulatory therapies and brain-machine interfaces (BMIs) that can facilitate a more regulated and predictable control over functionally relevant volumetric neurotransmission.

## Results

### A customized pipeline enables *in vivo* noradrenergic modulation and brain-wide single-nucleus RNA profiling

The LC-NE system is known to achieve a diverse set of functions in part through distinct modes of activity. Short-term periods of tonic LC hyperactivity are known to occur during events of high arousal or acute stress and can last for hours to days, resulting in physiological and behavioral changes. Similarly, LC-NE hypoactivity is observed in several pathologies, including narcolepsy, depression, and during a coma. To mimic such events and facilitate a comprehensive characterization of brain-wide transcriptional changes associated with LC-NE hyper- or hypoactivity, we used virus delivered Cre-dependent DREADDs to chemogenetically activate LC-NE neurons in *Dbh*^*cre*^ mice **(Figure 1a, b)**. We profiled tissue uniformly sampled from a single hemisphere across all brain regions in eight female mouse brains.

**Figure 1:**
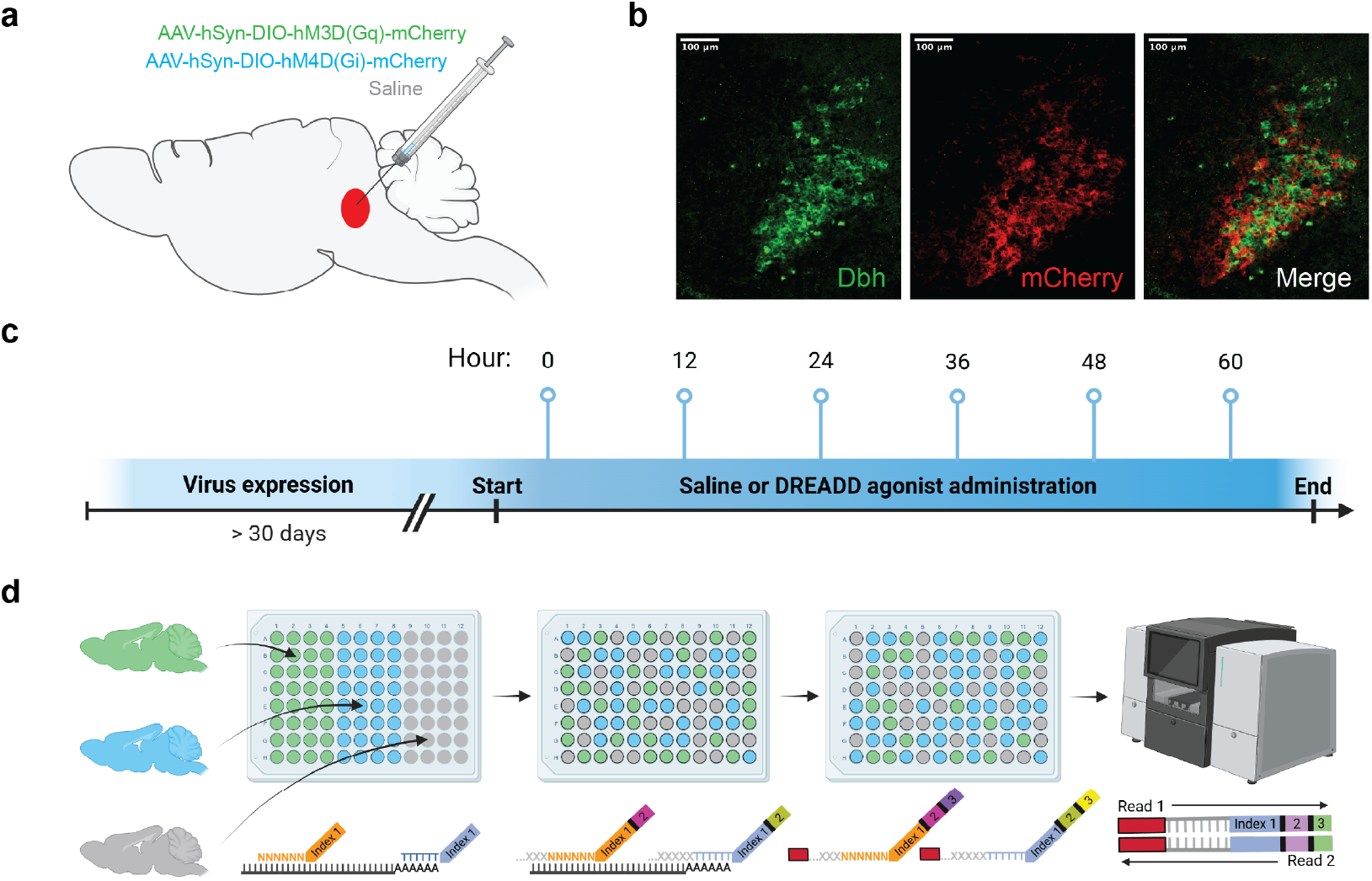
Experimental workflow and single-nucleus RNA sequencing library prep. **a)** Three groups of Dbh-Cre mice received bilateral LC injections of Gq-DREADD, Gi-DREADD, or saline. **b)** Representative histological images of the LC. **c)** Experimental timeline. **d)** EasySci combinatorial indexing library preparation.

To characterize brain-wide transcriptional changes following LC activation, mice were bilaterally injected with adeno associated viruses (AAVs) encoding Gq- or Gi-coupled DREADD targeting LC noradrenergic neurons and administered JHU37160 (a DREADD agonist) twice per day for three days prior to sacrifice **(Figure 1c**). snRNA-seq was performed on representative sections of the whole brain, resulting in the recovery of ∼55k high-quality nuclei across 8 samples **(Figure 1d)**. A UMAP clustering of nuclei revealed clean separation between classes according to the conventions of the Allen Brain Cell Atlas **(Figure 2a)**. Cell class identities were defined using hierarchical clustering and marker gene expression, capturing a broad diversity of neuronal and non-neuronal populations. Nuclei were annotated by transcriptomic class **(Figure 2b)**, major brain region **(Figure 2c)**, and neurotransmitter identity **(Figure 2d)**.

**Figure 2:**
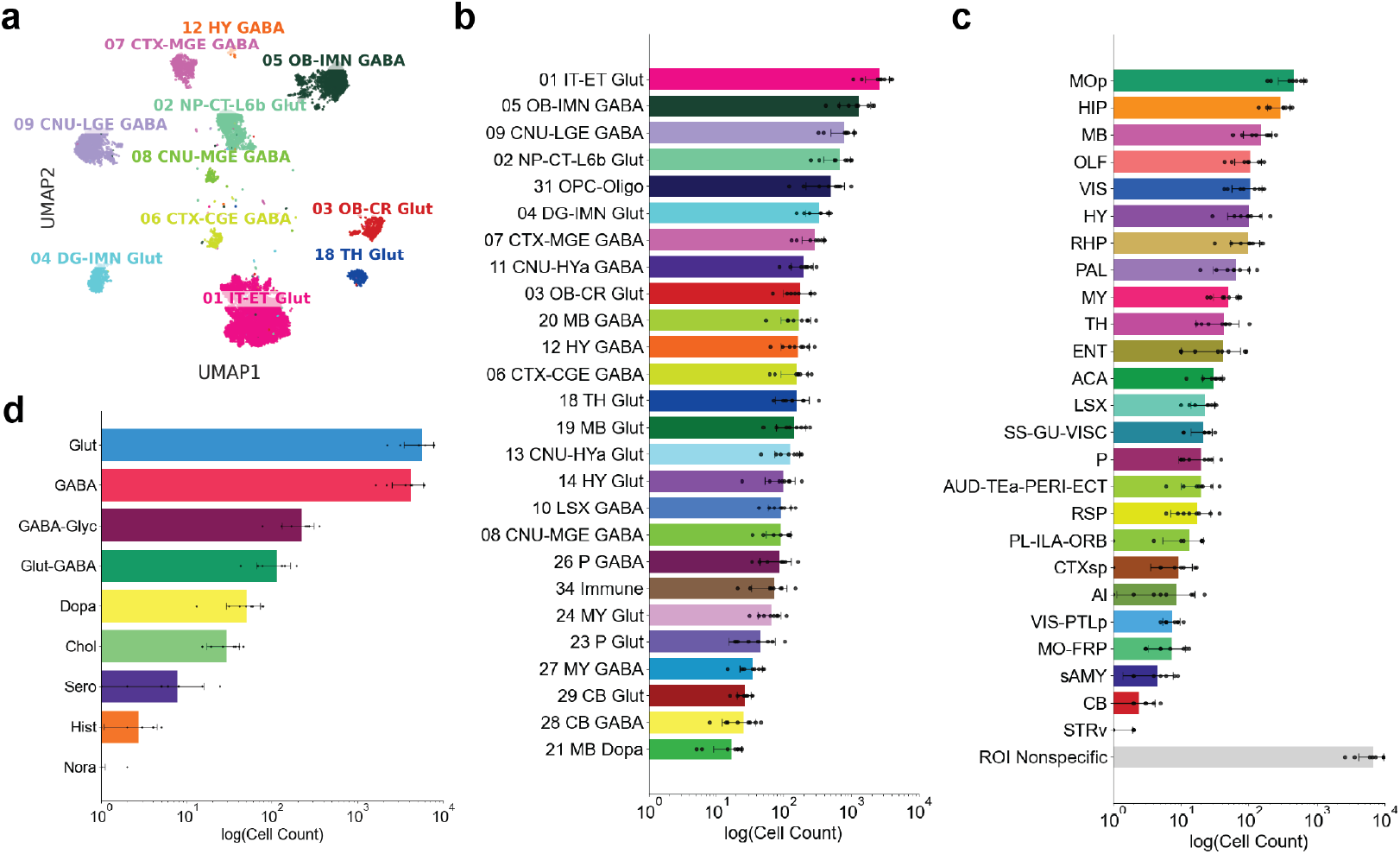
Clustering and composition of recovered nuclei. **a)** UMAP clustering labeled by cell type class. **b)** Composition of recovered nuclei by class, **c)** major brain region, and **d)** neurotransmitter type.

Nuclei were assessed on standard measures of quality control including the distribution of unique molecular identifiers (UMIs) per nucleus and the distribution of features per nucleus. Distributions were visualized across the neuronal and non-neuronal divide **(Figure 3a)**, treatments **(Figure 3b)**, classes **(Figure 3c)**, and brain regions **(Figure 3d)**.

**Figure 3:**
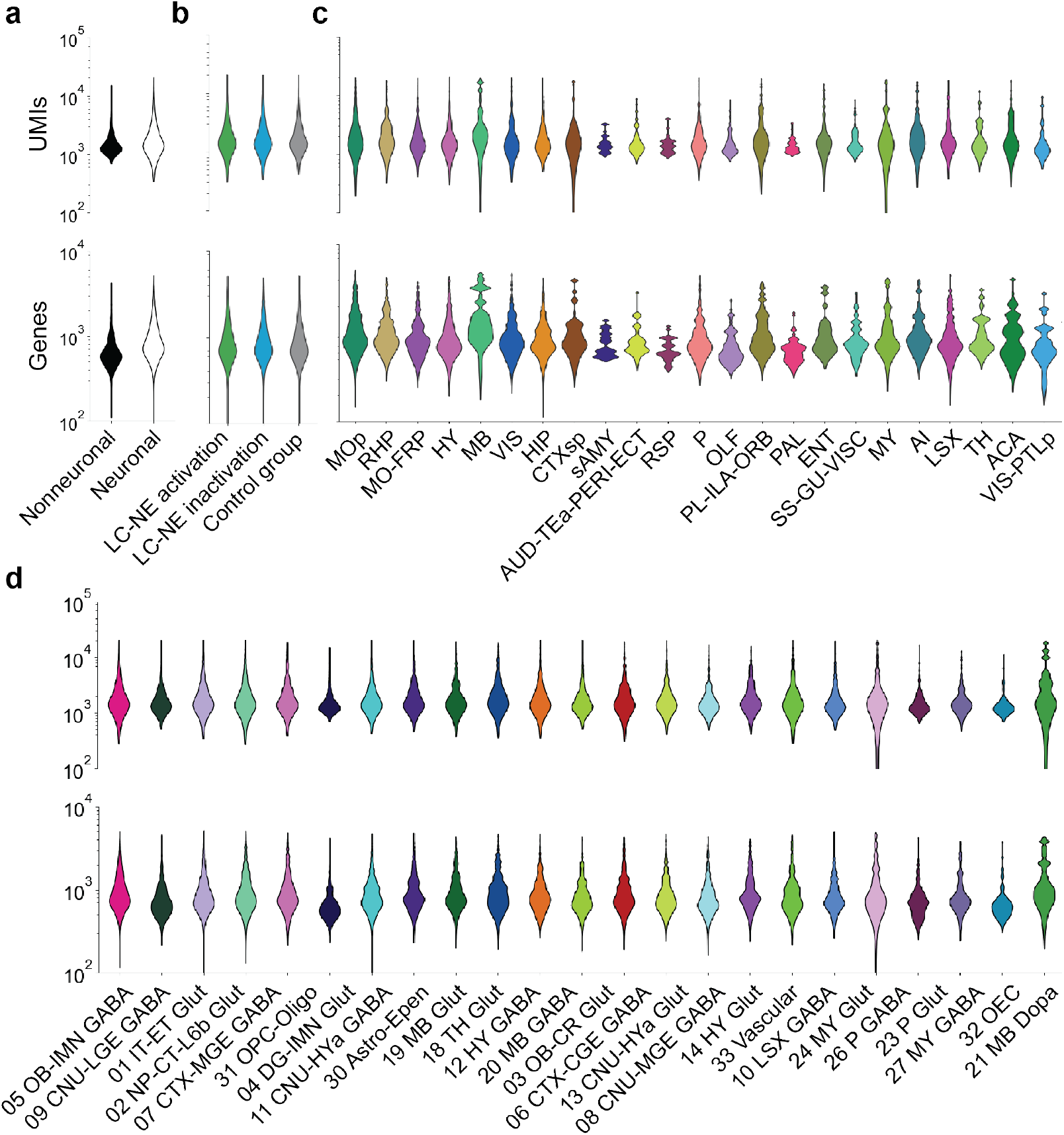
UMI and unique gene distributions. **a)** UMI and unique gene distributions between nuclei of nonneuronal and neuronal cells, **b)** treatment groups, **c)** major brain regions, and **d)** classes.

### Adrenergic receptor subtype expression is highly cell-type-specific

Expression levels of alpha- and beta-adrenergic receptors were quantified across classes **(Figure 4)**. Receptor expression patterns were heterogeneous, with classes expressing varied levels of *α*_1_, *α*_2_, or *β* receptors. In some classes, certain receptor subtypes were expressed at far lower amounts than other adrenergic receptor subtypes. For example, there is a distinct lack of the *α*_2*b*_, *α*_2*c*_, and *β*_2_ subtypes in the *30 Astro-Epen* class, whereas *01 IT-ET Glut* neurons express these adrenergic receptor subtypes. This finding suggests that different classes of cells exposed to the same levels of free norepinephrine may produce vastly different cellular outcomes.

**Figure 4:**
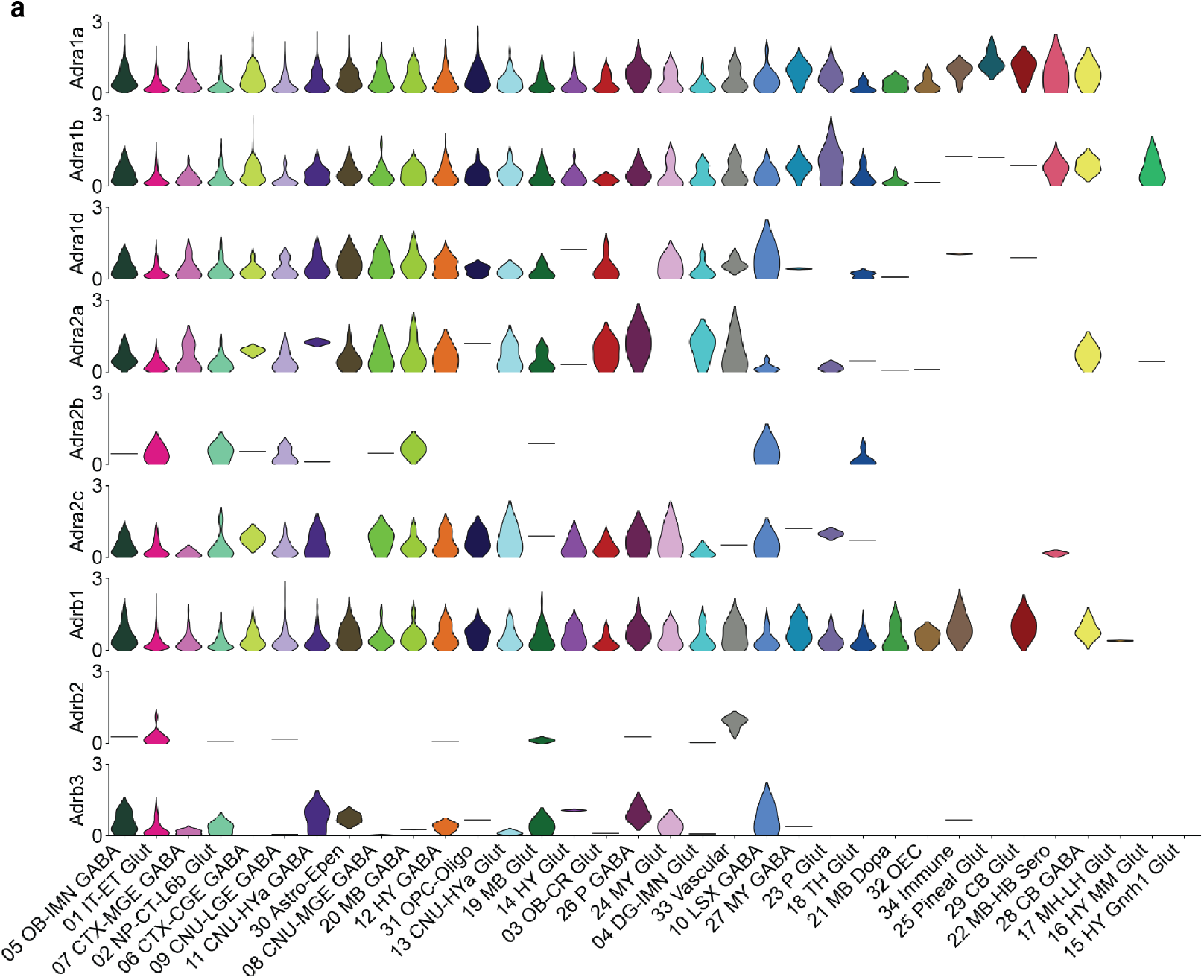
Adrenergic receptor subtype distribution across classes.

### Differential gene expression analysis reveals that LC activation affects the expression of more genes than LC inhibition

Differential gene expression analysis was performed to assess the impact of sustained locus coeruleus (LC) hyperactivity across annotated cell classes **(Figure 5a,b)**. Among all populations, three classes exhibited a relatively high number of differentially expressed genes in response to tonic LC hyperactivation.

**Figure 5:**
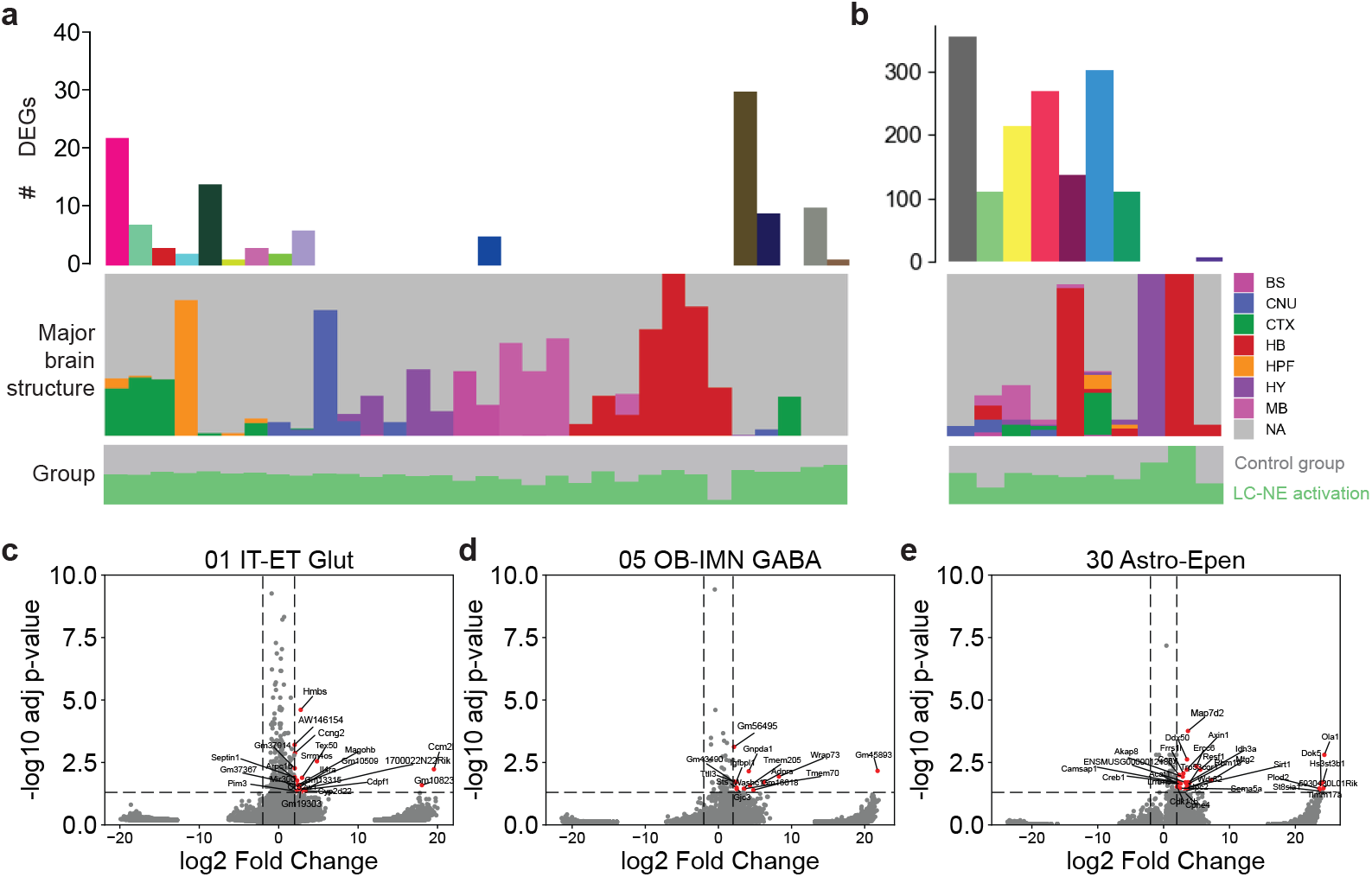
Differential gene expression summary, tonic LC hyperactivity vs. controls. **a)** DEG summary across classes. *Top:* Number of differentially expressed genes across classes. *Middle:* Breakdown of nuclei proportion by brain region across classes. *Bottom:* Breakdown of nuclei proportion by treatment group across classes. **b)** DEG summary across dominant neurotransmitter identities. *Top:* Number of differentially expressed genes across dominant neurotransmitter identities. *Middle:* Breakdown of nuclei proportion by brain region across dominant neurotransmitter identities. *Bottom:* Breakdown of nuclei proportion by treatment group across dominant neurotransmitter identities. **c-e)** Volcano plots for the top three classes by number of differentially expressed genes. Significance thresholds set to absolute log_2_ fold change > 2 and adj. p-value < 0.05.

In *01 IT-ET Glut* neurons, elevated expression of *Hmbs, Ccrn2*, and *Tex50* was observed, suggesting that long range intratelencephalic and extratelencephalic projection neurons undergo transcriptional alterations associated with metabolic and transcriptional regulation **(Figure 5c)**. In *05 OB-IMN GABA*neurons, which includes neurons of the olfactory bulb, increased expression of *Gm45893, Wrap73*, and *Gnpda1* was detected, consistent with shifts in RNA processing and amino sugar metabolism **(Figure 5d)**. In *30 Astro-Epen* glial population, encompassing astrocytes and ependymal cells, *Ola1, Map7d2*, and *Dok5* were among the genes upregulated, pointing to changes in cytoskeletal dynamics and MAPK signaling **(Figure 5e)**.

Comparatively fewer DEGs were identified from tonic LC hypoactivity analysis **(Figure 6a,b)**. In *01 IT-ET Glut* neurons, downregulation of *Eif4ebp1* and upregulation of *Gm24407*, and *Acr* was detected, indicating potential shifts in synaptic architecture **(Figure 6c)**. In *22 MB-HB Sero* neurons, a midbrain-hindbrain serotonergic class, increased expression of *Cpne7* and *Slc8a1* was observed, suggesting alterations in calcium handling and membrane excitability in neuromodulatory brainstem circuits **(Figure 6d)**. Within the *30 Astro Epen* population, decreased expression of *Prlr* and increased expression of *Pde10a* was identified, implicating effects on prolactin signaling and cyclic nucleotide metabolism in astroglial function **(Figure 6e)**.

**Figure 6:**
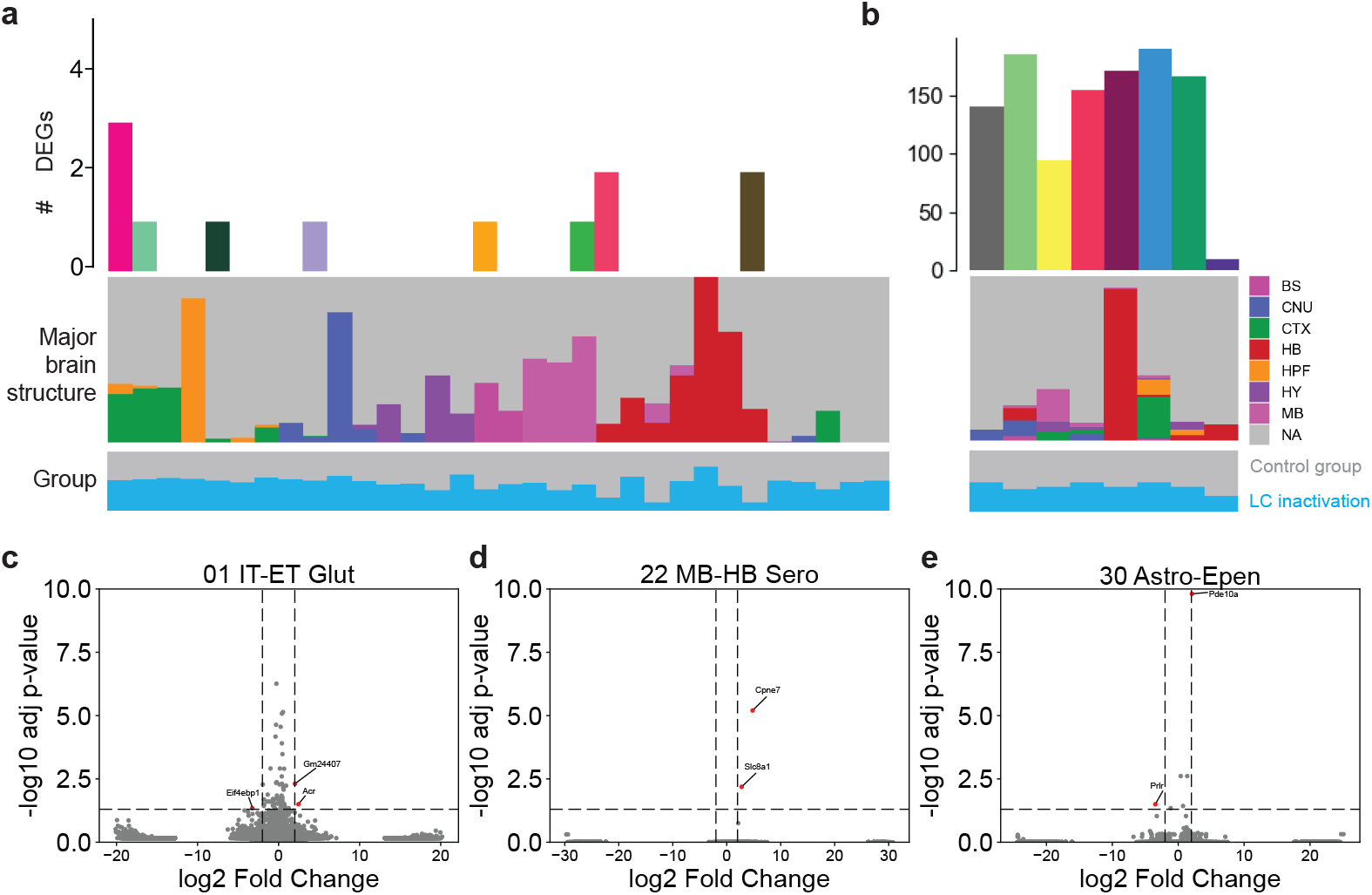
Differential gene expression summary, tonic LC hypoactivity vs. controls. **a)** DEG summary across classes. *Top:* Number of differentially expressed genes across classes. *Middle:* Breakdown of nuclei proportion by brain region across classes. *Bottom:* Breakdown of nuclei proportion by treatment group across classes. **b)** DEG summary across dominant neurotransmitter identities. *Top:* Number of differentially expressed genes across dominant neurotransmitter identities. *Middle:* Breakdown of nuclei proportion by brain region across dominant neurotransmitter identities. *Bottom:* Breakdown of nuclei proportion by treatment group across dominant neurotransmitter identities. **c-e)** Volcano plots for the top three classes by number of differentially expressed genes. Significance thresholds set to absolute log_2_ fold change > 2 and adj. p-value < 0.05.

These findings demonstrate that diminished tonic norepinephrine release leads to class-specific transcriptional adaptations, with differential sensitivity across neuronal and glial populations. Together, the data suggest that the absence of noradrenergic tone reconfigures homeostatic signaling pathways in a manner distinct from the gene programs engaged by LC hyperactivity.

### Gene set enrichment analysis reveals perturbation-specific functional programs

Gene set enrichment analysis revealed that tonic LC hyperactivity activated diverse molecular programs in a cell type-specific manner **(Figure 7)**. In *01 IT-ET Glut* neurons, enrichment was observed in pathways related to synaptic remodeling and neurotransmitter receptor organization. Notably, terms such as *presynaptic membrane organization* and *NMDA selective glutamate receptor signaling pathway* exhibited high enrichment scores, indicating that LC hyperactivity substantially modulates both synaptic architecture and receptor sensitivity in pyramidal neurons. In contrast, the *05 OB-IMN GABA* neuronal class was enriched for terms including regulation of *grooming behavior, positive regulation of the force of heart contraction*, and *regulation of heart rate by chemical signal*, suggesting a broader physiological role for these neurons of the olfactory bulb. Additionally, significantly decreased enrichment scores were identified for histone H3K27 methyltransferase activity across multiple cell types, pointing to a shared epigenetic response across classes to tonic LC hyperactivity.

**Figure 7:**
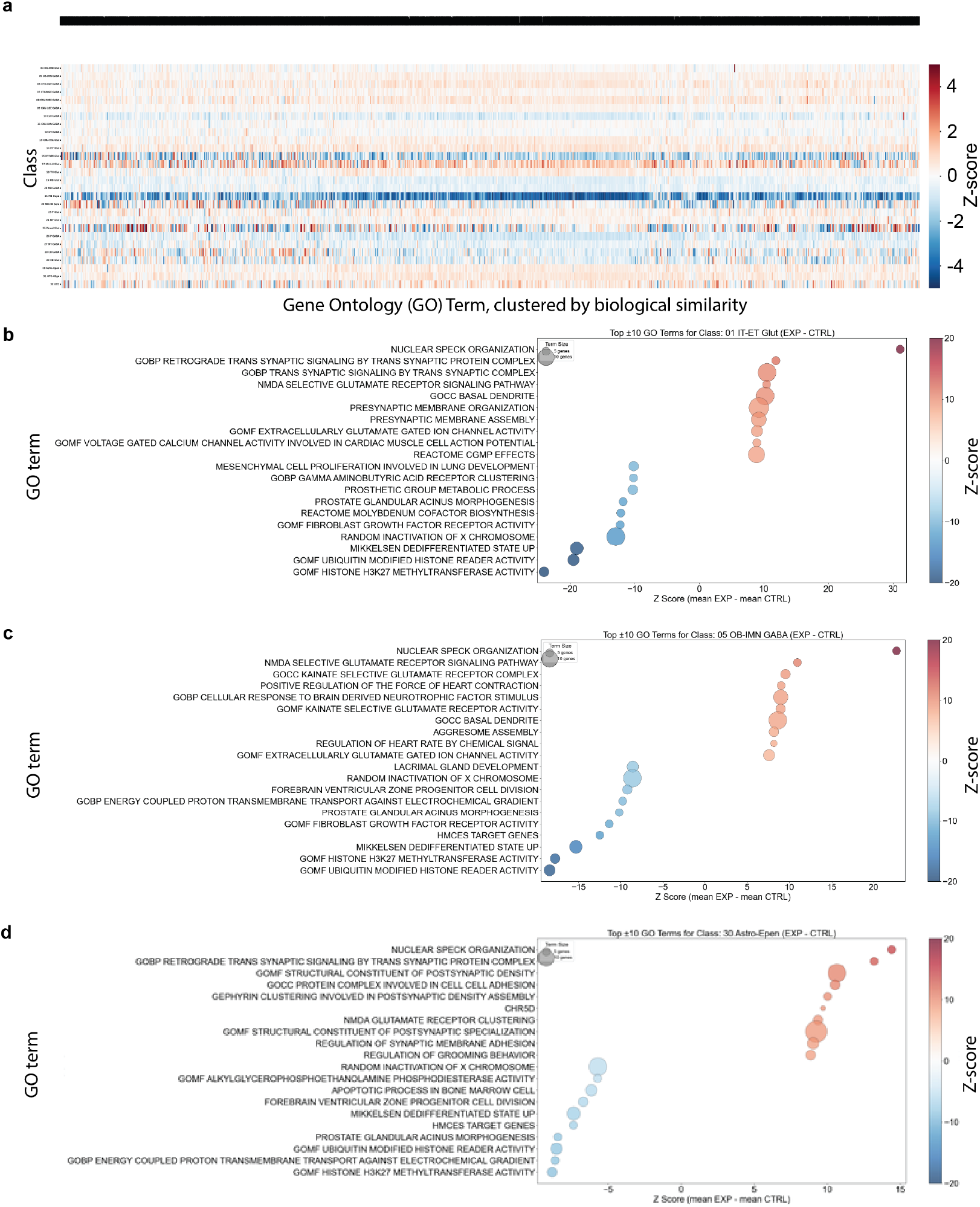
Gene set enrichment analysis, tonic LC hyperactivity vs. controls. **a)** Heatmap of classes (rows) vs. gene ontology (GO) terms (columns). **b-d)** Bubble plots of highly relevant GO terms for the top three classes by number of DEGs.

## Methods

### Animals

All procedures were approved by the Institutional Animal Care and Use Committee (IACUC) at Columbia University. B6.Cg-Dbh^tm3.2(cre)Pjen/J^(Dbh-Cre, JAX stock #033951) mice and C57BL/6J WT mice aged 8 weeks were used for all experiments. Mice were housed in a 12-hour light/dark cycle with *ad libitum* access to food and water.

### Stereotaxic surgery and viral injections

Aseptic surgical procedures for the injection of AAV viral vectors are similar to those described previously [15, 48]. Briefly, mice were anesthetized with isoflurane (5% induction, 1.5% maintenance) and subsequently secured in a stereotaxic frame (Kopf Instruments) with a heating pad (FHC, Bowdoinham, ME) to maintain body temperature. The skull was carefully leveled and fur above the scalp was clipped and then removed with depilatory cream. An alternating application of Betadine and 70% alcohol solution was used to disinfect the surgical site. Prior to incision, buprenorphine (0.05 mg/kg) and saline solution containing 5% dextrose (1 mL) were administered subcutaneously to provide analgesia and hydration for the duration of the surgery. A local anesthetic (lidocaine, 2%) was also applied to the incision site before a midline scalp incision was made.

For all AAV injections, burr holes were made above the LC region, and saline was applied to each craniotomy to prevent desiccation of exposed brain surfaces. Pulled capillary glass micropipettes (Drummond Scientific, Broommall, PA) were back-filled with AAV solution and injected into the target brain regions at a rate of 23 nL/sec using a precision injection system (Nanoliter 2020, World Precision Instruments, Sarasota, FL). The micropipette was left in place for at least 3 minutes following each injection and then slowly withdrawn. To selectively target LC neurons for chemogenetic manipulations, either AAV-hSyn-DIO-hM4D(Gi)-mCherry (Addgene Cat. Number: 44362) or AAV-hSyn-DIO-hM3D(Gq)-mCherry (Addgene Cat. Number: 44361) was injected into the LC of both hemispheres (300 nL at each injection site, AP: -5.4 mm, ML: +/-1.0 mm, DV from dura: 2.5 mm, 3.0 mm, and 3.5 mm). Controls were injected with sterile phosphate buffered saline (PBS). At the conclusion of the surgery, Baytril (5 mg/kg) and carprofen (5 mg/kg) were administered. Four additional doses of Baytril and two additional doses of carprofen were administered every 24 hours after surgery. Animals’ weights were measured at least once per day for 5 days.

### Chemogenetic perturbation

DREADD agonist JHU37160 (0.2 mg/kg; Hello Bio, Cat.#HB6261) dissolved in sterile saline was injected intraperitoneally into Dbh-Cre mice to inactivate hM4D(Gi)-expressing or activate hM3D(Gq)-expressing neurons in the target region. Saline of equivalent volume was administered as a control. Injections were given every 12 hours for 3 consecutive days, and each animal was euthanized through cervical dislocation 30 minutes following the final injection.

### Recovery of neural tissue

Immediately following euthanasia, neural tissue was extracted, split into hemispheres down the midline, and flash frozen in chilled isopentane (2-methylbutane). Each right hemisphere was allocated for nuclei isolation, and the left hemisphere was allocated for histology.

### Recovery and purification of nuclei for single-nucleus RNA-sequencing

For snRNAseq library preparation, brains were sagittaly sectioned at 10 *µ*m across the entirety of one hemisphere. Every 6th slice was collected in an Eppendorf tube for subsequent processing. Tissue was then dissociated and nuclei isolated using an optimized method [49], and combinatorial barcodes were added to each transcript following protocols established previously [50, 51].

### Library preparation

#### Histology

Frozen tissue sections were stored at –80°C until use. To begin staining, slides were equilibrated to room temperature and placed on a 60°C slide warmer for 5 minutes. A hydrophobic barrier was drawn around each section and allowed to dry for 10 minutes. Sections were rehydrated with two 5-minute washes in 1× PBS.

Antigen retrieval was performed by incubating the slides in 0.01 M sodium citrate buffer (pH 6.0) at 60°C for 30 minutes. The buffer was composed of 135 mL dH_2_O, 2.7 mL 0.1 M citric acid, and 12.3 mL 0.1 M sodium citrate. Following antigen retrieval, endogenous peroxidase activity was blocked by incubating slides in a solution of 10% methanol and 3% hydrogen peroxide in dH_2_O for 20 minutes at room temperature on a shaker (150 rpm). Proper chemical waste disposal protocols were followed for methanol-containing solutions. Slides were then gently blotted to remove excess liquid and incubated in a pre-treatment solution containing 1× PBS, 10% donkey serum, and 1% Triton X-100 for 30 minutes at room temperature to permeabilize tissues and reduce nonspecific binding.

For blocking of endogenous mouse IgG, sections were incubated for 2 hours in donkey anti-mouse IgG Fab fragments (1.3 mg/mL; diluted 1:10 in blocking buffer). This step was included for experiments involving fluorophores with emission in the 633–647 nm range. The Fab carrier solution consisted of 10% donkey serum, 1% Triton X-100 (from a 20% stock in PBS), and 1× PBS. Following blocking, slides were washed eight times for 4 minutes each: four times with PBS-T (PBS + 0.1% Tween-20) and four times with PBS alone (alternating between the two, with the final rinse in PBS only).

Slides were then incubated with primary antibodies diluted in 1°Ab carrier solution (10% donkey serum, 1% Triton X-100, and 1× PBS) for 48 hours at 4°C in a humidified chamber. Antibody solutions were centrifuged briefly before dilution and vortexed thoroughly to ensure mixing. Control slides were incubated with carrier solution lacking primary antibody.

On the second day, slides were washed three times for 5 minutes each in 1× PBS. Secondary antibody incubation was performed for 6 hours at room temperature in the dark, using antibodies diluted in PBS. After incubation, slides were again washed three times in 1× PBS in the dark. For mounting, sections were coverslipped using Fluoromount-G with or without DAPI and allowed to dry overnight in a dark, dry chamber at room temperature. Slides were sealed with clear nail polish (optional) and stored in the dark until imaging.

### Computational methods

#### Loading data and examining nuclei recovery

To begin the analysis, single-nucleus RNA sequencing data were loaded using `load_adata()`, a function that accounted for the operating system and local machine name. This ensured cross-platform compatibility and reproducibility of results. The resulting AnnData object served as the foundational data structure for all downstream analyses. To characterize the cellular composition of the dataset, nuclei were grouped according to transcriptionally defined cell classes derived from the Allen Brain Cell Atlas. The relative abundance of each class was computed on a per-sample basis, and the mean and standard deviation were visualized as horizontal bar plots with the `class_composition()` function. **(Figure 2B)**. This allowed for a comparative analysis of the recovery rates of various classes across samples and highlighted potential biases in nuclei recovery.

To assess regional representation, nuclei were further stratified by anatomical location using the `region_composition()`function. Only nuclei uniquely registered to a single region were considered. For each brain region, the average percentage across samples and its standard deviation were plotted, with per sample variability overlaid as scatter points **(Figure 2C)**. To account for nuclei assigned to multiple regions, the `region_composition()` function includes a parameter, `egistration_category=multi`, that allows for using expanded annotations from multi-mapped registrations, wherein each region was given equal weight. Additionally, nuclei that lacked spatial assignments were included as a separate category labeled ‘None’, ensuring complete representation using `region_composition_all_labels()`. This approach enabled a holistic view of anatomical sampling density and spatial registration fidelity.

Nuclei were then categorized by their dominant neurotransmitter phenotype with the `neurotransmitter_composition()` function. Sample-level percentages were computed for each neurotransmitter group, and their averages and standard deviations were visualized using horizontal bar plots **(Figure 2d)**. This analysis provided insight into the chemical diversity of the sampled population and facilitated downstream differential gene expression comparisons based on neurotransmitter identity.

#### Quality control

To evaluate data quality, the distributions of total UMI counts and detected gene features were visualized using `umi_feature_class(), umi_feature_treatment()`, and `umi_feature_region()`, which produced violin plots across different classes, experimental treatments, and brain regions, respectively **(Figure 3)**. For each panel, data were filtered to exclude extreme low- and high-UMI counts, thereby removing outliers and damaged nuclei. This QC step ensured that comparisons across samples, treatments, and classes reflected true biological variability rather than technical noise.

#### Cell type labeling

All recovered cells were assigned a hierarchical cell type label using the MapMyCells tool provided by the Allen Institute [44]. Cells with a bootstrapping probability less than 0.7 for class assignment were excluded from dataset.

#### Adrenergic receptor subtype expression

Expression of adrenergic receptor subtypes was visualized across multiple classes using `gene_expression_class()`, which generated violin plots, color-coded to remain consistent with earlier functions. **(Figure 4)**. It should be noted that this function can be used to probe transcriptional changes of any annotated gene in response to perturbation, at cell type resolution. All panels were restricted to cells expressing the gene of interest and used normalized expression values.

To summarize the number of differentially expressed genes per class **(Figures 5a, 6a)**, a function, `deg_class_summary()`, created. Within this function, Scanpy’s built-in `rank_genes_groups()` function was used on each class subset, and DEGs were filtered based on fold change and adjusted p-value thresholds. Classes with insufficient experimental or control nuclei were excluded. The number of DEGs per class was visualized as a bar plot, reflecting the magnitude of response per cell type. In addition, `deg_class_summary()` also generates volcano plots for the top three classes by number of DEGs to illustrate the spread and significance of individual genes **(Figures 5c, 6c)**. The distribution of nuclei by major brain structure within each class was visualized using a stacked bar plot using `region_proportion_class()`. Counts were aggregated and normalized by class to obtain region proportions, and regions were color-coded based on a predefined palette. This revealed spatial biases or enrichment patterns in nuclei distribution for each class. To determine whether observed transcriptional differences could be attributed to treatment-related differences in class composition, the proportions of nuclei per class were compared between treatment and control conditions with `treatment_proportion_class()`. Nuclei counts were normalized across groups, and class-level proportions were visualized using a dual-stacked bar chart. This facilitated the interpretation of DEG results in the context of potential compositional shifts due to noradrenergic modulation.

An analogous analysis was performed at the neurotransmitter level using `deg_neurotransmitter_summary(), region_proportion_neurotransmitter()`, and `treatment_proportion_neurotransmiter()` **(Figures 5b, 6b)**. The number of DEGs was computed for each neurotransmitter group using pseudo bulk profiles and Welch’s t-tests, mirroring the methodology used in class-based analysis. The regional composition of each neurotransmitter class was plotted to assess anatomical biases. The treatment composition per neurotransmitter group was visualized, enabling cross-comparison between chemical identity, spatial distribution, and treatment exposure.

#### Gene set enrichment analysis

To identify enriched biological processes, gene set enrichment analysis was performed using `go_heatmap_clustered()`, which precompiled gene sets from MSigDB and structured Gene Ontology (GO) terms from the Gene Ontology Consortium. For each class, expression differences between treatment and control conditions were computed and used to score gene sets based on mean expression shifts. Z-score normalization was applied, and the top most variable GO terms were selected. Gene sets were hierarchically clustered by Jaccard distance and visualized as a heatmap, revealing shared and divergent pathway-level responses across classes **(Figure 7a)**.

To probe class-specific gene ontology trends at higher resolution, GO enrichment was summarized with `go_class()` at the individual class level using bubble plots **(Figure 7b)**. For a selected class, gene sets with the highest and lowest Z-scores were displayed, with bubble size encoding the number of contributing genes and color encoding expression bias. Term descriptions were fetched via API queries to ensure interpretability. This analysis provided focused insight into the specific biological processes modulated by treatment in individual cell classes.

## Discussion

Using chemogenetic activation and inactivation of LC neurons and single-nucleus RNA sequencing their downstream molecular consequences, this study, for the first time, characterized brain-wide, cell type–specific transcriptional responses to repeated overactivation or inactivation of the LC. Our findings demonstrate that neuromodulatory perturbation exerts widespread but highly cell type- and region-specific gene expression changes **(Figures 5, 6b)**. These results highlight the capacity of a single neuromodulatory nucleus to enact broad, context-dependent molecular programs through volumetric signaling.

The observed effects reinforce the understanding that neuromodulators such as norepinephrine influence neural circuits not solely through point-to-point synaptic transmission but also via diffuse, extrasynaptic mechanisms [52]. This volumetric mode of transmission enables broad modulation of neuronal excitability, plasticity, gene expression, and function over biological scales from the molecular level to the organismal level. The present data suggest that the specific transcriptional outcomes of LC-NE signaling are shaped by the distribution of adrenergic receptor subtypes across cell classes and support a model in which the adrenergic receptor combinations and ratios of each cell type functions as a molecular gate, filtering global NE signals into discrete, cell-type-specific responses **(Figure 4)**.

It is notable that diverse functional programs were activated in a cell type-specific manner **(Figure 7)**. For example, in tonic LC hyperactivity, *01 IT-ET Glut* exhibited enrichment in pathways related to synaptic remodeling and neurotransmitter receptor organization. Indeed, GO terms like presynaptic membrane organization, NMDA selective glutamate receptor signaling pathway had especially high enrichment scores, suggesting that LC hyperactivity heavily adjusts both the connections and sensitivity of intratelencephalic and extratelencephalic glutamatergic neurons. Under the same treatment condition, cells belonging to the *05 OB-IMN GABA* class were enriched in genes relevant to the regulation of grooming behavior, positive regulation of the force of heart contraction, and regulation of heart rate by chemical signal, pointing to the potential influence of immature olfactory bulb neurons in broader behavior and physiology. Taken together, these results indicate that LC neuromodulation does not simply elicit a uniform molecular response but instead modulates functionally tailored transcriptional programs depending on the intrinsic identity and role of the target cells. Notably, however, this study also identified significantly lowered enrichment scores for histone H3K27 methyltransferase activity across at least three classes, illustrating the paradigm’s ability to also capture commonalities of molecular response to neural perturbation across cell types.

The effects of LC perturbation observed in this study likely reflect a mixture of rapid, reversible signaling cascades and slower gene regulatory programs. While brief episodes of LC activation have been associated with short-term changes in synaptic dynamics and network activity during sensory and cognitive processing [11, 53], the present paradigm lays the foundation for future inquiries into prolonged neuromodulatory tone shifts. Longitudinal investigations of LC modulation are relevant to pathophysiological states in which chronic LC hypoactivity (e.g., in narcolepsy, depression) or hyperactivity (e.g., in anxiety disorders or neurodegeneration) is observed. The LC-NE system is also essential for normal neural functions such as learning, cognition, circadian rhythm entrainment, and sleep in behavior [30, 54, 55, 56]; the gene expression changes identified here may therefore provide a molecular framework for understanding how persistent alterations in neuromodulatory tone contribute to both the etiology of these pathophysiological states as well as the mechanisms of neural functions at baseline.

The transcriptomic framework established in this study is broadly generalizable and can be applied to the investigation of other neuromodulatory systems. The combination of whole-brain snRNA-seq with cell-type-specific annotations allows for systematic mapping of the molecular consequences of neural perturbations with high spatial and cellular resolution. This approach provides a scalable method to evaluate how global neurotransmitter fluctuations are decoded across the brain and offers a foundation for the development of transcriptionally-informed interventions.

Several future investigations are warranted based on the findings of this hypothesis-generating study. First, although we observed significant gene expression changes induced by LC perturbation, the perturbation in the present study was relatively short-term, consisting of twice-daily treatments over three days. However, the pathogenesis of many neuropsychiatric disorders often involves chronic hyperactivation or hypoactivation of the LC-NE system [57]. Therefore, a brain-wide characterization of transcriptomic changes across diverse cell types in response to long-term LC-NE perturbation, which mimicks pathogenesis of neuropsychiatric disorders such as chronic stress, could yield critical insights into the molecular mechanisms underlying these disorders. Second, the integration of behavioral and physiological measurements would allow for the connection of transcriptional changes to functional outcomes. For instance, concurrent assessment of behavior, neural dynamics (e.g., calcium imaging, electrophysiology), and gene expression during LC perturbation would enable causal inference regarding the impact of neuromodulatory tone cell-type-specific gene expression on perception, learning, or decision making. Such multimodal approaches would be essential for linking observations across biological scales, from molecular adaptations to systems-level functions and behavior. Finally, while the current study employed a binary model of neuromodulatory modulation (hyperactive vs. hypoactive), future studies could explore graded or temporally dynamic perturbations to better approximate endogenous LC activity patterns. Such studies would be instrumental in elucidating how the brain decodes continuous fluctuations in neuromodulator levels as well as the nature of nonlinear relationships between neuromodulatory tone, gene regulatory outcomes, and higher-order behavior.

Together, this work demonstrates that defined neuromodulatory perturbations give rise to widespread, cell type-specific transcriptional changes across the brain. These findings advance our understanding of the molecular mechanisms underlying volumetric neuromodulation and establish a platform for the development of next-generation, transcriptionally informed neuromodulatory therapies and brain-machine interfaces [58, 59, 60]. By framing gene expression as a programmable substrate of neuromodulatory control, this study lays the groundwork for engineering more predictable and targeted interventions that engage brain functions across biological and temporal scales relevant to both neural restoration and enhancement.

## Disclaimer

Q.W. is the co-founder of Sharper Sense. T.L. owns stock options in Envisagenics from his previous employment.

## References

[1] Thomas J. Brozoski, Roger M. Brown, H. E. Rosvold, and Patricia S. Goldman. Cognitive Deficit Caused by Regional Depletion of Dopamine in Prefrontal Cortex of Rhesus Monkey. Science, 205(4409):929–932, August 1979. Publisher: American Association for the Advancement of Science.

[2] S. G. Birnbaum, P. X. Yuan, M. Wang, S. Vijayraghavan, A. K. Bloom, D. J. Davis, K. T. Gobeske, J. D. Sweatt, H. K. Manji, and A. F. T. Arnsten. Protein kinase C overactivity impairs prefrontal cortical regulation of working memory. Science (New York, N.Y.), 306(5697):882–884, October 2004.

[3] Eve Marder. Neuromodulation of Neuronal Circuits: Back to the Future. Neuron, 76(1):1–11, October 2012. Publisher: Elsevier.

[4] Susan J. Sara. The locus coeruleus and noradrenergic modulation of cognition. Nature Reviews. Neuroscience, 10(3):211–223, March 2009.

[5] Seung-Hee Lee and Yang Dan. Neuromodulation of Brain States. Neuron, 76(1):209–222, October 2012. Publisher: Elsevier.

[6] Angela J. Yu and Peter Dayan. Uncertainty, Neuromodulation, and Attention. Neuron, 46(4):681–692, May 2005. Publisher: Elsevier.

[7] Stefano Fusi, Wael F. Asaad, Earl K. Miller, and Xiao-Jing Wang. A Neural Circuit Model of Flexible Sensorimotor Mapping: Learning and Forgetting on Multiple Timescales. Neuron, 54(2):319–333, April 2007. Publisher: Elsevier.

[8] Michael C. Avery and Jeffrey L. Krichmar. Neuromodulatory Systems and Their Interactions: A Review of Models, Theories, and Experiments. Frontiers in Neural Circuits, 11:108, December 2017.

[9] Paul Greengard. The Neurobiology of Slow Synaptic Transmission. Science, 294(5544):1024–1030, November 2001. Publisher: American Association for the Advancement of Science.

[10] Brian J. Schriver, Svetlana Bagdasarov, and Qi Wang. Pupil-linked arousal modulates behavior in rats performing a whisker deflection direction discrimination task. Journal of Neurophysiology, 120(4):1655–1670, October 2018. Publisher: American Physiological Society.

[11] Charles Rodenkirch, Yang Liu, Brian J. Schriver, and Qi Wang. Locus coeruleus activation enhances thalamic feature selectivity via norepinephrine regulation of intrathalamic circuit dynamics. Nature Neuroscience, 22(1):120– 133, January 2019. Publisher: Nature Publishing Group.

[12] Farzan Nadim and Dirk Bucher. Neuromodulation of Neurons and Synapses. Current opinion in neurobiology, 0:48–56, December 2014.

[13] Andrea Francesca Salvador, Kalil Alves de Lima, and Jonathan Kipnis. Neuromodulation by the immune system: a focus on cytokines. Nature Reviews Immunology, 21(8):526–541, August 2021. Publisher: Nature Publishing Group.

[14] Silvia Graciela Ruginsk, Andre de Souza Mecawi, Melina Pires da Silva, Wagner Luis Reis, Ricardo Coletti, Juliana Bezerra Medeiros de Lima, Lucila Leico Kagohara Elias, and Jose Antunes-Rodrigues. Gaseous Modulators in the Control of the Hypothalamic Neurohypophyseal System. Physiology, 30(2):127–138, March 2015. Publisher: American Physiological Society.

[15] Yuxiang (Andy) Liu, Yuhan Nong, Jiesi Feng, Guochuan Li, Paul Sajda, Yulong Li, and Qi Wang. Phase synchrony between prefrontal noradrenergic and cholinergic signals indexes inhibitory control, January 2025. Pages: 2024.05.17.594562 Section: New Results.

[16] Lindsay A. Schwarz, Kazunari Miyamichi, Xiaojing J. Gao, Kevin T. Beier, Brandon Weissbourd, Katherine E. DeLoach, Jing Ren, Sandy Ibanes, Robert C. Malenka, Eric J. Kremer, and Liqun Luo. Viral-genetic tracing of the input–output organization of a central noradrenaline circuit. Nature, 524(7563):88–92, August 2015. Publisher: Nature Publishing Group.

[17] Liqun Luo. Architectures of neuronal circuits. Science, 373(6559):eabg7285, September 2021. Publisher: American Association for the Advancement of Science.

[18] Özge D. Özçete, Aditi Banerjee, and Pascal S. Kaeser. Mechanisms of neuromodulatory volume transmission. Molecular Psychiatry, 29(11):3680–3693, November 2024. Publisher: Nature Publishing Group.

[19] Ben Tsuda, Stefan C. Pate, Kay M. Tye, Hava T. Siegelmann, and Terrence J. Sejnowski. Neuromodulators generate multiple context-relevant behaviors in a recurrent neural network by shifting activity flows in hyperchannels, September 2024. Pages: 2021.05.31.446462 Section: New Results.

[20] C Nicholson, GT Bruggencate, R Steinberg, and H Stöckle. Calcium modulation in brain extracellular microenvironment demonstrated with ion-selective micropipette. Proceedings of the National Academy of Sciences, 74(3):1287–1290, March 1977. Publisher: Proceedings of the National Academy of Sciences.

[21] He J. V. Zheng, Qi Wang, and Garrett B. Stanley. Adaptive shaping of cortical response selectivity in the vibrissa pathway. Journal of Neurophysiology, 113(10):3850–3865, June 2015. Publisher: American Physiological Society.

[22] L. F. Agnati, M. Zoli, I. Strömberg, and K. Fuxe. Intercellular communication in the brain: Wiring versus volume transmission. Neuroscience, 69(3):711–726, December 1995.

[23] L. F. Agnati, G. Leo, A. Zanardi, S. Genedani, A. Rivera, K. Fuxe, and D. Guidolin. Volume transmission and wiring transmission from cellular to molecular networks: history and perspectives. Acta Physiologica, 187(1-2):329–344, 2006. _eprint: https://onlinelibrary.wiley.com/doi/pdf/10.1111/j.1748-1716.2006.01579.x.

[24] David A. McCormick, Dennis B. Nestvogel, and Biyu J. He. Neuromodulation of Brain State and Behavior. Annual Review of Neuroscience, 43(Volume 43, 2020):391–415, July 2020. Publisher: Annual Reviews.

[25] Edward Zagha and David A McCormick. Neural control of brain state. Current Opinion in Neurobiology, 29:178–186, December 2014.

[26] Yue Liang, Wu Shi, Anfeng Xiang, Dandan Hu, Liecheng Wang, and Ling Zhang. The NAergic locus coeruleus ventrolateral preoptic area neural circuit mediates rapid arousal from sleep. Current Biology, 31(17):3729–3742.e5, September 2021. Publisher: Elsevier.

[27] Hanna Hayat, Noa Regev, Noa Matosevich, Anna Sales, Elena Paredes-Rodriguez, Aaron J. Krom, Lottem Bergman, Yong Li, Marina Lavigne, Eric J. Kremer, Ofer Yizhar, Anthony E. Pickering, and Yuval Nir. Locus coeruleus norepinephrine activity mediates sensory-evoked awakenings from sleep. Science Advances, 6(15):eaaz4232, April 2020. Publisher: American Association for the Advancement of Science.

[28] Brandon R. Munn, Eli J. Müller, Gabriel Wainstein, and James M. Shine. The ascending arousal system shapes neural dynamics to mediate awareness of cognitive states. Nature Communications, 12(1):6016, October 2021. Publisher: Nature Publishing Group.

[29] Lindsay A. Schwarz and Liqun Luo. Organization of the Locus Coeruleus-Norepinephrine System. Current Biology, 25(21):R1051–R1056, November 2015.

[30] Gina R. Poe, Stephen Foote, Oxana Eschenko, Joshua P. Johansen, Sebastien Bouret, Gary Aston-Jones, Carolyn W. Harley, Denise Manahan-Vaughan, David Weinshenker, Rita Valentino, Craig Berridge, Daniel J. Chandler, Barry Waterhouse, and Susan J. Sara. Locus coeruleus: a new look at the blue spot. Nature Reviews Neuroscience, 21(11):644–659, November 2020. Publisher: Nature Publishing Group.

[31] Kimberly L. Simpson, Daniel W. Altman, Li Wang, Michael L. Kirifides, Rick C.-S. Lin, and Barry D. Waterhouse. Lateralization and functional organization of the locus coeruleus projection to the trigeminal somatosensory pathway in rat. Journal of Comparative Neurology, 385(1):135–147, 1997. _eprint: https://onlinelibrary.wiley.com/doi/pdf/10.1002/%28SICI%291096-9861%2819970818%29385%3A1%3C135%3A%3AAID-CNE8%3E3.0.CO%3B2-3.

[32] Richard H. Masland. Neuronal cell types. Current Biology, 14(13):R497–R500, July 2004. Publisher: Elsevier.

[33] Melanie Philipp and Lutz Hein. Adrenergic receptor knockout mice: distinct functions of 9 receptor subtypes. Pharmacology & Therapeutics, 101(1):65–74, January 2004.

[34] Orit Barrett and Talya Wolak. 23 - Peripheral Adrenergic Blockers. In George L. Bakris and Matthew J. Sorrentino, editors, Hypertension: A Companion to Braunwald’s Heart Disease (Third Edition), pages 222–229. Elsevier, January 2018.

[35] Dianne M. Perez. The Adrenergic Receptors: In the 21st Century. Springer Science & Business Media, October 2007. Google-Books-ID: ffdGnMIoZYQC.

[36] Dianne M. Perez. 1-Adrenergic Receptors in Neurotransmission, Synaptic Plasticity, and Cognition. Frontiers in Pharmacology, 11, September 2020. Publisher: Frontiers.

[37] T. C. Westfall. Sympathomimetic Drugs and Adrenergic Receptor Antagonists. In Larry R. Squire, editor, Encyclopedia of Neuroscience, pages 685–695. Academic Press, Oxford, January 2009.

[38] Eduardo Listik and Qin Wang. Alpha2-adrenergic receptors. In Italo Biaggioni, Kirsteen Browning, Gregory Fink, Jens Jordan, Phillip A. Low, and Julian F. R. Paton, editors, Primer on the Autonomic Nervous System (Fourth Edition), pages 49–52. Academic Press, January 2023.

[39] Masahiro Kaneko and Tomoyuki Takahashi. Presynaptic Mechanism Underlying cAMP-Dependent Synaptic Potentiation. The Journal of Neuroscience, 24(22):5202–5208, June 2004.

[40] Wagner Steuer Costa, Szi-chieh Yu, Jana F. Liewald, and Alexander Gottschalk. Fast cAMP Modulation of Neurotransmission via Neuropeptide Signals and Vesicle Loading. Current Biology, 27(4):495–507, February 2017. Publisher: Elsevier.

[41] Brian P. Ramos and Amy F. T. Arnsten. Adrenergic pharmacology and cognition: Focus on the prefrontal cortex. Pharmacology & Therapeutics, 113(3):523–536, March 2007.

[42] Xinyu Xu, Jonas Kaindl, Mary J. Clark, Harald Hübner, Kunio Hirata, Roger K. Sunahara, Peter Gmeiner, Brian K. Kobilka, and Xiangyu Liu. Binding pathway determines norepinephrine selectivity for the human 1AR over 2AR. Cell Research, 31(5):569–579, May 2021.

[43] Julius T. Dongdem, Axandrah E. Etornam, Solomon Beletaa, Issah Alidu, Hassan Kotey, and Cletus A. Wezena. The 3-Adrenergic Receptor: Structure, Physiopathology of Disease, and Emerging Therapeutic Potential. Advances in Pharmacological and Pharmaceutical Sciences, 2024(1):2005589, 2024. _eprint: https://onlinelibrary.wiley.com/doi/pdf/10.1155/2024/2005589.

[44] Allen Brain Cell Atlas.

[45] Zizhen Yao, Cindy T. J. van Velthoven, Michael Kunst, Meng Zhang, Delissa McMillen, Changkyu Lee, Won Jung, Jeff Goldy, Aliya Abdelhak, Matthew Aitken, Katherine Baker, Pamela Baker, Eliza Barkan, Darren Bertagnolli, Ashwin Bhandiwad, Cameron Bielstein, Prajal Bishwakarma, Jazmin Campos, Daniel Carey, Tamara Casper, Anish Bhaswanth Chakka, Rushil Chakrabarty, Sakshi Chavan, Min Chen, Michael Clark, Jennie Close, Kirsten Crichton, Scott Daniel, Peter DiValentin, Tim Dolbeare, Lauren Ellingwood, Elysha Fiabane, Timothy Fliss, James Gee, James Gerstenberger, Alexandra Glandon, Jessica Gloe, Joshua Gould, James Gray, Nathan Guilford, Junitta Guzman, Daniel Hirschstein, Windy Ho, Marcus Hooper, Mike Huang, Madie Hupp, Kelly Jin, Matthew Kroll, Kanan Lathia, Arielle Leon, Su Li, Brian Long, Zach Madigan, Jessica Malloy, Jocelin Malone, Zoe Maltzer, Naomi Martin, Rachel McCue, Ryan McGinty, Nicholas Mei, Jose Melchor, Emma Meyerdierks, Tyler Mollenkopf, Skyler Moonsman, Thuc Nghi Nguyen, Sven Otto, Trangthanh Pham, Christine Rimorin, Augustin Ruiz, Raymond Sanchez, Lane Sawyer, Nadiya Shapovalova, Noah Shepard, Cliff Slaughterbeck, Josef Sulc, Michael Tieu, Amy Torkelson, Herman Tung, Nasmil Valera Cuevas, Shane Vance, Katherine Wadhwani, Katelyn Ward, Boaz Levi, Colin Farrell, Rob Young, Brian Staats, Ming-Qiang Michael Wang, Carol L. Thompson, Shoaib Mufti, Chelsea M. Pagan, Lauren Kruse, Nick Dee, Susan M. Sunkin, Luke Esposito, Michael J. Hawrylycz, Jack Waters, Lydia Ng, Kimberly Smith, Bosiljka Tasic, Xiaowei Zhuang, and Hongkui Zeng. A high-resolution transcriptomic and spatial atlas of cell types in the whole mouse brain. Nature, 624(7991):317–332, December 2023. Publisher: Nature Publishing Group.

[46] Lei Han, Zhen Liu, Zehua Jing, Yuxuan Liu, Yujie Peng, Huizhong Chang, Junjie Lei, Kexin Wang, Yuanfang Xu, Wei Liu, Zihan Wu, Qian Li, Xiaoxue Shi, Mingyuan Zheng, He Wang, Juan Deng, Yanqing Zhong, Hailin Pan, Junkai Lin, Ruiyi Zhang, Yu Chen, Jinhua Wu, Mingrui Xu, Biyu Ren, Mengnan Cheng, Qian Yu, Xinxiang Song, Yanbing Lu, Yuanchun Tang, Nini Yuan, Suhong Sun, Yingjie An, Wenqun Ding, Xing Sun, Yanrong Wei, Shuzhen Zhang, Yannong Dou, Yun Zhao, Luyao Han, Qianhua Zhu, Junfeng Xu, Shiwen Wang, Dan Wang, Yinqi Bai, Yikai Liang, Yuan Liu, Mengni Chen, Chun Xie, Binshi Bo, Mei Li, Xinyan Zhang, Wang Ting, Zhenhua Chen, Jiao Fang, Shuting Li, Yujia Jiang, Xing Tan, Guolong Zuo, Yue Xie, Huanhuan Li, Quyuan Tao, Yan Li, Jianfeng Liu, Yuyang Liu, Mingkun Hao, Jingjing Wang, Huiying Wen, Jiabing Liu, Yizhen Yan, Hui Zhang, Yifan Sheng, Shui Yu, Xiaoyan Liao, Xuyin Jiang, Guangling Wang, Huanlin Liu, Congcong Wang, Ning Feng, Xin Liu, Kailong Ma, Xiangjie Xu, Tianyue Han, Huateng Cao, Huiwen Zheng, Yadong Chen, Haorong Lu, Zixian Yu, Jinsong Zhang, Bo Wang, Zhifeng Wang, Qing Xie, Shanshan Pan, Chuanyu Liu, Chan Xu, Luman Cui, Yuxiang Li, Shiping Liu, Sha Liao, Ao Chen, Qing-Feng Wu, Jian Wang, Zhiyong Liu, Yidi Sun, Jan Mulder, Huanming Yang, Xiaofei Wang, Chao Li, Jianhua Yao, Xun Xu, Longqi Liu, Zhiming Shen, Wu Wei, and Yan-Gang Sun. Single-cell spatial transcriptomic atlas of the whole mouse brain. Neuron, 0(0), March 2025. Publisher: Elsevier.

[47] Meng Zhang, Xingjie Pan, Won Jung, Aaron R. Halpern, Stephen W. Eichhorn, Zhiyun Lei, Limor Cohen, Kimberly A. Smith, Bosiljka Tasic, Zizhen Yao, Hongkui Zeng, and Xiaowei Zhuang. Molecularly defined and spatially resolved cell atlas of the whole mouse brain. Nature, 624(7991):343–354, December 2023. Publisher: Nature Publishing Group.

[48] Aditya Apte, Julia Fernald, Cody Slater, Marc Sorrentino, Brett Youngerman, and Qi Wang. Bidirectional Modulation of Somatostatin-expressing Interneurons in the Basolateral Amygdala Reduces Neuropathic Pain Perception in Mice, April 2025. Pages: 2025.03.28.645947 Section: New Results.

[49] Beth K. Martin, Chengxiang Qiu, Eva Nichols, Melissa Phung, Rula Green-Gladden, Sanjay Srivatsan, Ronnie Blecher-Gonen, Brian J. Beliveau, Cole Trapnell, Junyue Cao, and Jay Shendure. Optimized single nucleus transcriptional profiling by combinatorial indexing. Nature protocols, 18(1):188, January 2023. Publisher: NIH Public Access.

[50] Andras Sziraki, Ziyu Lu, Jasper Lee, Gabor Banyai, Sonya Anderson, Abdulraouf Abdulraouf, Eli Metzner, Andrew Liao, Jason Banfelder, Alexander Epstein, Chloe Schaefer, Zihan Xu, Zehao Zhang, Li Gan, Peter T. Nelson, Wei Zhou, and Junyue Cao. A global view of aging and Alzheimer’s pathogenesis-associated cell population dynamics and molecular signatures in human and mouse brains. Nature Genetics, 55(12):2104–2116, December 2023. Publisher: Nature Publishing Group.

[51] Sanjay R. Srivatsan, José L. McFaline-Figueroa, Vijay Ramani, Lauren Saunders, Junyue Cao, Jonathan Packer, Hannah A. Pliner, Dana L. Jackson, Riza M. Daza, Lena Christiansen, Fan Zhang, Frank Steemers, Jay Shendure, and Cole Trapnell. Massively multiplex chemical transcriptomics at single-cell resolution. Science, 367(6473):45– 51, January 2020. Publisher: American Association for the Advancement of Science.

[52] Virginia M. Pickel. Extrasynaptic distribution of monoamine transporters and receptors. In Progress in Brain Research, volume 125 of Volume Transmission Revisited, pages 267–276. Elsevier, January 2000.

[53] Ernesto Durán, Mingyu Yang, Ricardo Neves, Nikos K. Logothetis, and Oxana Eschenko. Modulation of Prefrontal Cortex Slow Oscillations by Phasic Activation of the Locus Coeruleus. Neuroscience, 453:268–279, January 2021.

[54] Oxana Eschenko, Cesare Magri, Stefano Panzeri, and Susan J. Sara. Noradrenergic neurons of the locus coeruleus are phase locked to cortical up-down states during sleep. Cerebral Cortex (New York, N.Y.: 1991), 22(2):426–435, February 2012.

[55] Alejandro Osorio-Forero, Georgios Foustoukos, Romain Cardis, Najma Cherrad, Christiane Devenoges, Laura M. J. Fernandez, and Anita Lüthi. Infraslow noradrenergic locus coeruleus activity fluctuations are gatekeepers of the NREM–REM sleep cycle. Nature Neuroscience, 28(1):84–96, January 2025. Publisher: Nature Publishing Group.

[56] Celia Kjaerby, Mie Andersen, Natalie Hauglund, Verena Untiet, Camilla Dall, Björn Sigurdsson, Fengfei Ding, Jiesi Feng, Yulong Li, Pia Weikop, Hajime Hirase, and Maiken Nedergaard. Memory-enhancing properties of sleep depend on the oscillatory amplitude of norepinephrine. Nature Neuroscience, 25(8):1059–1070, August 2022. Publisher: Nature Publishing Group.

[57] Jennifer A. Ross and Elisabeth J. Van Bockstaele. The Locus Coeruleus-Norepinephrine System in Stress and Arousal: Unraveling Historical, Current, and Future Perspectives. Frontiers in Psychiatry, 11, January 2021. Publisher: Frontiers.

[58] Michael Jigo, Jason B. Carmel, Qi Wang, and Charles Rodenkirch. Transcutaneous cervical vagus nerve stimulation improves sensory performance in humans: a randomized controlled crossover pilot study. Scientific Reports, 14(1):3975, February 2024. Publisher: Nature Publishing Group.

[59] Charles Rodenkirch and Qi Wang. Rapid and transient enhancement of thalamic information transmission induced by vagus nerve stimulation. Journal of Neural Engineering, 17(2):026027, April 2020. Publisher: IOP Publishing.

[60] Charles Rodenkirch, Jason B. Carmel, and Qi Wang. Rapid Effects of Vagus Nerve Stimulation on Sensory Processing Through Activation of Neuromodulatory Systems. Frontiers in Neuroscience, 16, July 2022. Publisher: Frontiers.

